# ProfPPIdb: pairs of physical protein-protein interactions predicted for entire proteomes

**DOI:** 10.1101/332510

**Authors:** Linh Tran, Tobias Hamp, Burkhard Rost

## Abstract

**Motivation:** Protein-protein interactions (PPIs) play a key role in many cellular processes. Most annotations of PPIs mix experimental and computational data. The mix optimizes coverage, but obfuscates the annotation origin. Some resources excel at focusing on reliable experimental data. Here, we focused on new pairs of interacting proteins for several model organisms based solely on sequence-based prediction methods.

**Results:** We extracted reliable experimental data about which proteins interact (binary) for eight diverse model organisms from public databases, namely from *Escherichia coli, Schizosaccharomyces pombe, Plasmodium falciparum, Drosophila melanogaster, Caenorhabditis elegans, Mus musculus, Rattus norvegicus, Arabidopsis thaliana*, and for the previously used *Homo sapiens* and *Saccharomyces cerevisiae*. Those data were the base to develop a PPI prediction method for each model organism. The method used evolutionary information through a profile-kernel Support Vector Machine (SVM). With the resulting eight models, we predicted all possible protein pairs in each organism and made the top predictions available through a web application. Almost all of the PPIs made available were predicted between proteins that have not been observed in any interaction, in particular for less well-studied organisms. Thus, our work complements existing resources and is particularly helpful for designing experiments because of its uniqueness. Experimental annotations and computational predictions are strongly influenced by the fact that some proteins have many partners and others few. To optimize machine learning, recent methods explicitly ignored such a network-structure and rely either on domain knowledge or sequence-only methods. Our approach is independent of domain-knowledge and leverages evolutionary information. The database interface representing our results is accessible from https://rostlab.org/services/ppipair/. The data can also be downloaded from https://figshare.com/collections/ProfPPI-DB/4141784.

## Introduction

### Operational definition of physical Protein-Protein Interactions (PPIs)

We define <u>PPIs</u> as interactions that bring two different proteins A and B directly into ‘physical contact’. This ‘molecular’ perspective on PPIs differs from the most frequent view of both associations and permanent complexes. For us the crucial aspect of a PPI is that it brings two proteins into direct physical contact (usually transiently, i.e. for a limited time). Given all PPIs in an organism, the *interactome* comprises all PPIs in the entire proteome; this network contains all non-temporal aspects of associations on the network level.

### Experimental annotations of binary PPI maps

Due to the importance, many experiments establish PPIs. Despite this effort, most pairs of physically interacting proteins remain likely unknown [1]. Statistical models of PPIs can amend the coverage of networks formed from binary PPIs (A binds B) cost-effectively by enriching protein association networks [2–4] or by combining heterogeneous data sources in Bayesian networks [5].

### Predictions important but often over-estimated

Numerous computational methods have been developed to predict protein-protein interactions using different data sources, e.g. secondary structure, phylogenetic tree, phylogenetic profile, and gene expression [6–10]. Most methods employ more than one of the mentioned properties. However, their application is limited due to their specific need of domain knowledge. These specific knowledge is but not universally available, and limit these methods to specific (smaller) datasets.

Further, many methods only use sequence information, such as motifs of co-occurrence on the level of domains [11–13], matching features from protein sequence, structure and evolutionary conservation for binding sites alone [10,14] and for binding sites and sequence/structure triads [15]. However, none of those sequence-based methods restrict their method to the identification of physical non-permanent PPIs as we defined them. Most of those methods used permanent complexes, the others also associations. This is also true for methods pioneering the use of kernel-based predictions [14,15]. Evolutionary information embedded in proteins sequence was employed to improve predicting PPIs [10,14,16,17], some in combination with profile kernels [18], by leveraging information available to us which are not domain specific.

Another set of problems with existing methods pertain to the problems in choosing “negatives”, i.e. pairs of proteins known not to interact [19]. In fact, negatives have to be carefully considered when setting up the cross-validation process [20]. Moreover, the cross-validation protocol also needs to carefully avoid using the same proteins in training and testing [21,22], and even allowing for homologues between training and testing over-estimates performance [20]. Overall, it appears that every careful independent review of existing methods has unraveled some substantial over-estimates [20–22]. One recent method combining profile kernels with Support Vector Machines (SVM) to predict pairs of physical, non-permanent PPIs has tried to avoid all known flaws [23]. However, it still awaits critical assessment from independent experts. This method improved particularly for proteins without experimental annotations about their interactions recommending the approach for discovery of novel PPIs [23].

Here, we simply apply the concept of profile-kernel SVMs [23] to the prediction of the entire interactomes in eight model organisms, namely ordered by size: *Escherichia coli, Schizosaccharomyces pombe* (fission yeast), *Plasmodium falciparum, Drosophila melanogaster* (fruit fly), *Caenorhabditis elegans* (roundworm), *Mus musculus* (mouse), *Rattus norvegicus* (rat), and *Arabidopsis thaliana* (mouse-ear cress). The choice of applying profile-kernel SVMs is due to its independence of domain knowledge and its usage of evolutionary profiles. Further, in vast evaluation we chose negative interactions by avoiding using the similar proteins in training and testing. Repeated cross-validation was employed to reduce additional over-estimation as stated in [20–22]. We have created a database of the most reliable predictions for each organism, and implemented a versatile online search interface (https://rostlab.org/services/ppipair/). Our new methods and new predictions at least double the number of organisms for which sequence-based PPI predictions are available, and they do this in a more consistent way than other method [24]. On top, our resource contributes the first-ever predictions for many un-annotated proteins.

## Materials and Methods

### Data Sources

We extracted PPIs from the following databases BioGRID [25], DIP [26], and IntAct [27]. BioGRID is a public curated database that holds 553,827 physical interactions from 58 species. DIP archives 795,534 PPIs from 777 organisms, curated both manually by experts and through computational approaches. IntAct is also public archiving 356,806 PPIs mostly from eight organisms. All PPIs originated either from publications or submissions from experimentalists.

### Data Extraction

We only used PPIs for which their protein identifiers mapped to the EBI reference proteomes [28]. We mapped proteins of each organism to a corresponding reference protein only if their sequences aligned with at least 95 % sequence identity. The fraction of PPIs that could not be mapped in this simple manner accounted for about 9 % of all data. We grouped the resulting PPIs by organism using taxonomy identifiers and differentiated PPIs from 768 organisms.

To predict PPIs, we needed as much reliable training data as possible. However, we also need to remove redundancy in many non-trivial ways [23]. We used an established expert knowledge-scoring scheme [29] to reflect the quality of evidence for a given PPI. The scheme assigned scores from one (lowest reliability) to ten (highest reliability) for each experimental method used to annotate a PPI. High scores were assigned to techniques such as *X-ray crystallography* or *electron tomography*, average scores of five were given to, e.g. complementation-based assays and affinity-based technologies. Methodologies that do not directly provide evidence for interaction, such as co-localization or co-sedimentation, were scored lowest. The scores are available online at our service. We applied that scheme to our PPI data and kept only PPIs with at least one experimental evidence ≥ 5. For instance, the *Escherichia coli* PPI between P0ABB0 and P0ABB4 is supported by two experimental methods: blue native page (score = 3) and pull down (score = 2.5); both below 5, i.e. we discarded this PPI. In contrast the PPI between P0ACF0 and P03004 established by enzyme linked immunosorbent assay (score = 5) was kept. After data filtering, we redundancy reduced the PPI set of each organism set such that no PPI pair was sequence-similar. A PPI pair was considered similar if at least one of the two sequences reached HVAL > 20 [30] to any protein already in the data set. Note that HVAL > 20 corresponds to > 40 % pairwise sequence identity for alignments over 250 residues.

We applied the above procedure to all 768 organisms for which we found PPIs. Only 8 of the 768 had at least 200 PPIs with strong experimental support. We considered these our ‘model organisms’. 200 PPI was the minimum number of data points we assumed to be necessary to train our method. Redundancy reduction shrank our data by over ten-fold for some organisms (Table 1). The most extreme attrition was for fly for which we extracted almost 80k PPIs from the databases, and could use only about 1.6k for training/testing.

**Table 1.** Data sets extracted from BioGRID, DIP and IntAct. <u>Organism</u>: latin name for eight model organisms sorted alphabetically; <u>NPPIs</u>: number of distinct physical pairs of protein-protein interactions extracted by merging the entire BioGRID, DIP, and IntAct; <u>NPPIs with strong evidence</u>: subset of previous column with reliable experimental evidence (according to [29]); <u>NPPIs with strong evidence redundancy reduced</u>: subset of previous column after removing sequence-similar pairs (HVAL > 20).

| Organism | NPPIs | NPPIs with strong evidence | NPPIs with strong evidence redundancy reduced |
| --- | --- | --- | --- |
| *A. thaliana* | 38,258 | 8,459 | 814 |
| *C. elegans* | 23,105 | 5,229 | 818 |
| *D. melanogaster* | 79,291 | 19,033 | 1,680 |
| *E. coli* | 27,119 | 8,587 | 998 |
| *M. musculus* | 30,070 | 6,262 | 734 |
| *P. falciparum* | 4,792 | 1,312 | 239 |
| *S. pombe* | 13,478 | 4,396 | 410 |
| *R. norvegicus* | 6,698 | 1,574 | 236 |
| *Sum over all 8* | 222,811 | 54,852 | 5,929 |

### Negative interactions

Databases collect positives (A binds B), i.e. PPIs with experimental evidence. For training, we also needed negatives (A does not bind B). We collected negatives as described before in [20], [23]. For each PPI data set, we sampled negatives in a ratio of 1:10 (10 negative for each positive). The 1:10 ratio seemed appropriate to provide enough negatives to sample the reality in a cell. As before in [20] and [23], we obtained negatives by randomly sampling from all possible combinations of proteins of an organism with the restrictions that each protein in a ‘negative PPI’ needed to differ in sequence (HVAL < 20) to all proteins in the positive training set.

### Profile-kernel SVM parameter optimization and cross-validation

Many advanced sequence-based PPI prediction methods have been developed. Park and Marcotte [22] showed that PIPE2 [24], Autocorrelation [31], and SigProd [32] performed well compared to other methods. We showed a profile-kernel SVM to improve over these methods for human and yeast [23]. This method is described in detail in [23]. The basic concept is described in the following. Essentially, the profile-kernel finds k-mers of k adjacent residues for which the conservation within a given protein family exceeds some value *σ* and then collects the most informative such k-mers through SVMs. Thus, as for each profile-kernel SVM [33], we needed to optimize two hyperparameters: the k-mer length k and the evolutionary score threshold *σ*. Following our previous experience, we sampled k = 3, 4, 5, 6 for *σ* = 4,…, 11. For all organisms with more than 500 non-redundant PPIs, we optimized the two parameters empirically with a grid-search on two-thirds of the PPI data for each organism (*training set*). The remaining third of each data set (*test set*) was used to confirm generalization. Each training set was split further into five parts. For every hyperparameter combination, we performed a full 5-fold cross-validation using four splits for *cross-validation training* and one for *cross-validation testing*. In this way, each of the five splits of the full non-redundant set was used as cross-training split exactly once. We repeated each 5-fold cross-validation five times from the start, including splitting positives and sampling negatives, in order to minimize sampling noise [34]. Finally, we used the best combination of k and *σ* and the entire training set to train the method one last time in order to predict the test set. For organisms with more than 200, but fewer than 500 PPIs (Table 1), we did not optimize parameters, but only evaluated their performance in a five times 5-fold cross-validation on the whole data set. As hyperparameters, we used the most common combination found for the larger PPI sets (*k* = 5, *σ* = 11).

### Evolutionary profiles

The evolutionary profiles were taken from PredictProtein [35]. They were created by PSI-BLAST-ing [36] queries against an 80 % non-redundant database combining UniProt [37] and PDB [38]. Our method never used any information not available through these profiles.

### Recall-precision curves

Each model built from a training data set outputs a score for each prediction. We used these scores to calculate precision-recall-curves. In a cross-validation, we used all precisions at a particular recall to calculate the mean and the standard deviation of the precisions at that point. If only one curve was available (assessment of hold-out sets for organisms with > 200 PPIs), we assumed precision to follow a standard binomial distribution and calculated the precision error at a particular recall as:

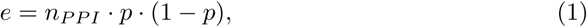
where *n_PPI_* denotes the number and p denotes the precision at that particular recall. In order to assess a particular parameter combination, we needed to condense the associated recall-precision curve into a single point. We did this by collecting all mean precision values until a recall of 20 % and then averaging over those values. The best parameter combination optimized this average precision.

### Interactome predictions

For predicting the entire interactomes, we used all available PPI data (training + test set) our models. As the hyper-parameters values *k* = 5, *σ* =11 yielded best performance for almost all organisms, we used those parameters for our interactome model for all organisms. This might not be the optimal solution, but it might provide the most conservative result avoiding more over-fitting. We applied our method to all pairs of proteins for which both proteins were dissimilar to any protein in the positives used for training. We chose to only publish the most reliable PPIs accounting to about 10 PPIs per protein of an organism (numbers given in Table 2).

**Table 2.** Whole interactome predictions. For each organism investigated, we aggregated the data we used for training and testing, trained a final model and predicted the whole interactome of that organism. <u>Organism</u>: latin name for eight model organisms sorted alphabetically; <u>Nprot</u>: number of proteins in proteome (values taken from [28]); <u>NpredPPI</u>: subset of PPIs used for prediction in which both proteins are dissimilar to the proteins in the positive interactions of the training set; <u>NprotPred</u>: corresponding number of proteins for which NpredPPI interactions were predicted, see Eq. 1 for calculation; <u>NpredPPI novel</u>: denotes the number of predicted PPIs for which both proteins are dissimilar to any known positive interaction, including redundant and low-quality PPIs; <u>NpredPPI ProfPPIdb</u>: subset with strongest predictions of previous column contained in our resource; <u>Nprot ProfPPIdb</u>: number of unique proteins in the PPIs published at https://rostlab.org/services/ppipair/, as well as https://figshare.com/collections/ProfPPI-DB/4141784.

| Organism | Nprot | NpredPPI<br>(NprotPred) | NpredPPI<br>novel | NpredPPI<br>ProfPPIdb | Nprot<br>ProtPPIdb |
| --- | --- | --- | --- | --- | --- |
| *A. thaliana* | 27,064 | 206,441,040<br>(20,320) | 71,251,953 | 250,000 | 7,023 |
| *C. elegans* | 20,137 | 142,171,953<br>(16,863) | 83,301,778 | 200,000 | 7,041 |
| *D. melanogaster* | 13,707 | 31,916,055<br>(7,990) | 5,410,405 | 100,000 | 7,664 |
| *E. coli* | 4,306 | 2,729,616<br>(2,337) | 332,520 | 40,000 | 1,341 |
| *M. musculus* | 22,136 | 131,325,321<br>(16,207) | 58,790,746 | 200,000 | 4,144 |
| *P. falciparum* | 5,159 | 9,041,878<br>(4,253) | 5,622,981 | 50,000 | 3,576 |
| *S. pombe* | 5,121 | 8,349,741<br>(4,087) | 2,630,071 | 50,000 | 3,946 |
| *R. norvegicus* | 21,330 | 211,922,578<br>(20,588) | 174,929,160 | 200,000 | 9,265 |
| *Sum over all 8* | 118,960 | 743,898,182<br>(92,645) | 402,269,614 | 1,090,000 | 44,000 |

## Results and Discussion

### Similar prediction performances between many organisms

Accumulating all non-redundant PPIs from the curated databases BioGRID, DIP and IntAct with reliable experimental annotations left only five organisms with over 500 PPIs enough to develop and evaluate organism-specific new methods using profile-kernel SVMs to predict PPIs from sequence: *Escherichia coli, Drosophila melanogaster, Caenorhabditis elegans, Mus musculus*, and *Arabidopsis thaliana* (Table 1). For each organism, two thirds of the data served for training and one-third as an independent test set. Training revealed that a *k*-mer length of *k* = 5 and conservation threshold *σ* = 11 were optimal for all organisms except *Escherichia coli* (Fig. 4). For simplicity, we used this hyper-parameter combination for all species (Fig. 1). Three other organisms *(Schizosaccharomyces pombe* (fission yeast), *Plasmodium falciparum*, and *Rattus norvegicus* (rat)) have too few experimental PPIs to fully optimize all parameters (Table 1: 236–410 PPIs). We evaluated the performance for these organisms in a 5-fold cross-validation using the default parameters *k* = 5, *σ* = 11 as fixed parameters (Fig. 2).

**Fig 1.**
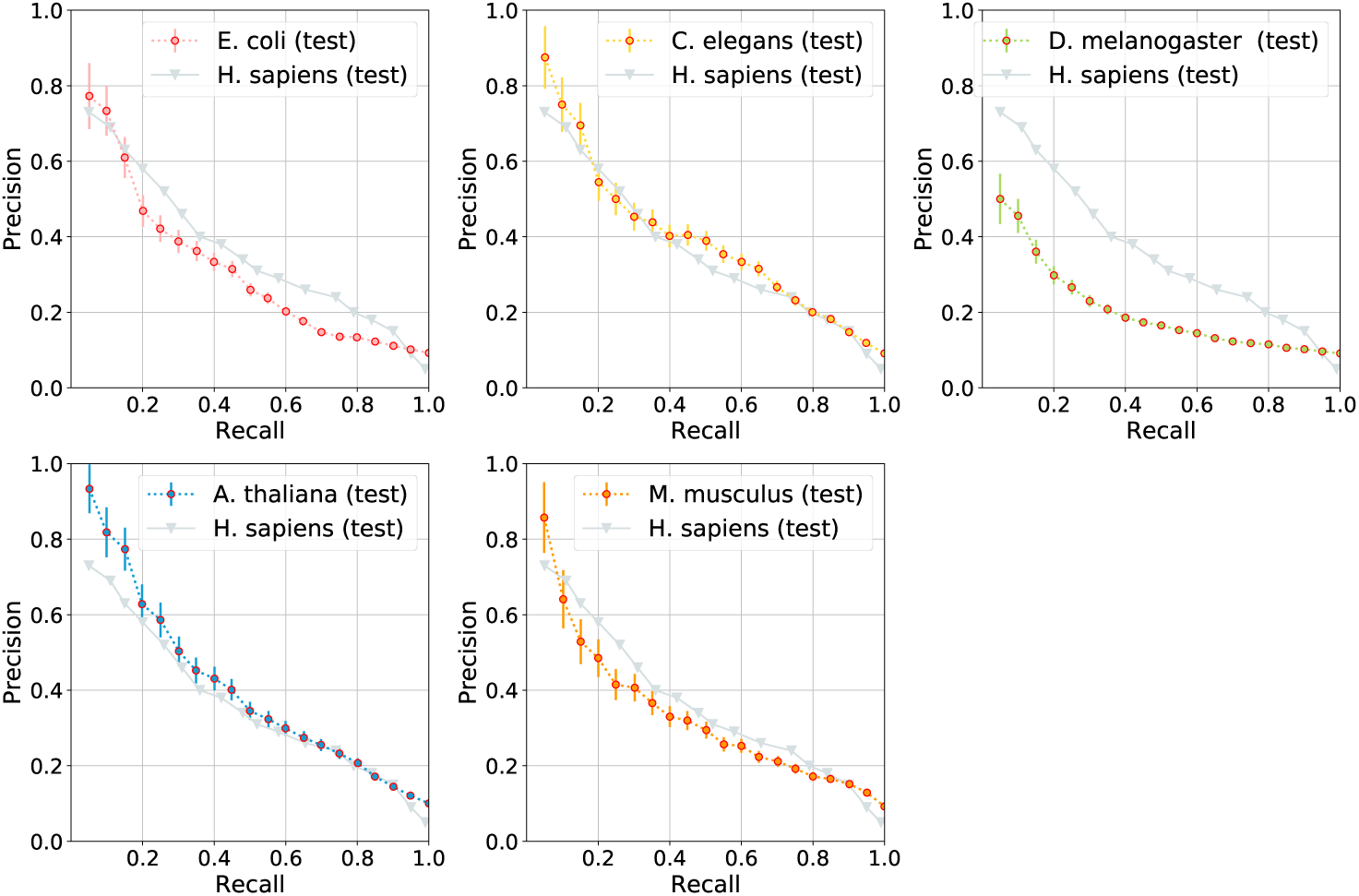
PPI test set for five organisms with ≥ 500 PPIs performed similar to human. The y-axes give precision (number of PPIs correctly predicted at threshold), the x-axes the recall (number of experimental interactions predicted at that threshold). The precision-recall curves of each organism describe the performance of the test data set. The model for that was trained with two-thirds of the PPI data. Bars give the standard binomial deviation; negatives were sampled at a rate of 10:1 (ten negatives for one positive). The gray values compare the model organisms to the PPI prediction performance for human. *H. sapiens* (test) denotes the performance of the same method described here for human PPIs.

**Fig 2.**
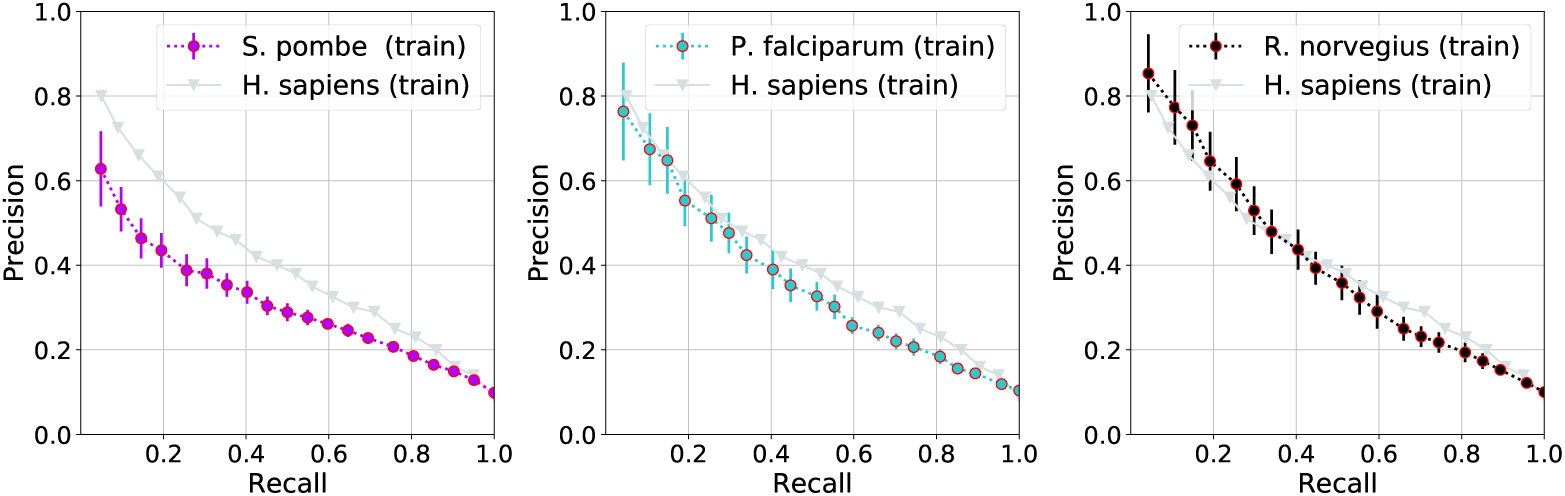
PPI test set for three organisms with < 500 PPIs inferior to human. The y-axes give precision (number of PPIs correctly predicted at threshold), the x-axes the recall (number of experimental interactions predicted at that threshold). The precision-recall curves of each organism describe the performance of the 5×5 cross validation of train data set. Bars give the standard deviation; negatives were sampled at a rate of 10:1 (ten negatives for one positive). The gray values compare the model organisms to the PPI prediction performance for human. *H. sapiens* (train) denotes the results of cross validation set.

For three of the five organisms (*Caenorhabditis elegans, Arabidopsis thaliana* and *Mus musculus*) our method performed on a similar level as our method predicting PPIs in human (Fig. 1). For low recall (≤ 0.1), the average precision for those three organisms appeared to even slightly (and significantly) exceed the values for human. However, our newly developed models for *Escherichia coli* and *Drosophila melanogaster* performed less well than the method for human. For *Escherichia coli*, changing the hyperparameters to *k* = 3, *σ* = 4 improved the performance (Fig. 5). We used the same hyperparameters for all eight models although we knew before using the testing set that this solution was not optimal. We did this as an additional precaution against over-fitting. For *Drosophila melanogaster* (fly) with over 1600 PPIs, we had no explanation for the dip in performance. In fact, the PPI predictions for fly appeared to be the worst amongst all ten organisms for which we applied our formalism (including human and baker’s yeast) although we had the highest number of PPIs for training. For fly we also observed by far the highest attrition from PPIs with 'some experimental evidence’ to ‘non-redundant PPIs with strong experimental evidence’ (Table 1: column ‘Number of PPIs’ vs. column ‘Number of PPIs with strong evidence’). However, we see no reason why this attrition should impact the consistency of the PPI data left over.

For organism with fewer than 500 PPIs (*Schizosaccharomyces pombe, Plasmodium falciparum* and *Rattus norvegicus*), we only evaluated the model performance with 5-fold cross-validation (Fig. 2). Our PPI prediction model for human appeared to perform better than the prediction models for these three organisms. This was most likely due to a lack of training data.

### Experimental evidences of novel predictions

We analysed our novel predictions by searching for any experimental evidence in databases such as BioGRID [25], DIP [26], IntAct [27], STRING [39], MINT [40] and Mentha [41]. All these databases have aggregated information of PPIs with experimental evidences. STRING [39], MINT [40] and Mentha [41] also provide confidence measures. Although the databases BioGRID [25], DIP [26], and IntAct [27] were already used for our organism-specific models, only a small subset of the databases’ PPIs was employed for training. The PPIs published on our online service only include PPIs which have not any experimental evidence from any of these three databases. In order to perform an evaluation of the quality of the predictions, we used the top 1 % of all predictions (ranked according to our confidence measure) which were not included in the training set. We compared these predictions against all experimental from BioGRID [25], DIP [26], and IntAct [27]. Overall, we found a total number of 772 PPIs with evidence which results in an average 86.79 % accuracy of correctly predicted PPIs. We also found evidences of PPIs for PPIs which our models did not predict any direct physical interaction. However, these evidences were usually experimental evidences with expert knowledge scores of lower or equal 4 [29] and thus highly likely to be false positives. A more detailed description of our findings can be found in the supplementary materials (Section A.2).

While we have only found minor number of PPIs with evidences in MINT [40] and Mentha [41], we found a significant portion of evidence in the STRING [39] database. Table 3 shows the number of evidences found of our evaluation with the STRING [39] database and includes numbers of evidences conforming with our predictions as well as the resulting accuracy. Except for *Mus musculus, Plasmodium falciparum* and *Rattus norvegicus*, we have found more than 1000 PPIs per organism with evidence in the database. With a high number of correctly predicted PPIs (both our prediction and STRING score indicate a PPI), we can observe a correlation between our most reliable PPIs and STRING PPI score. The average accuracy of positive predicted PPIs with STRING evidences is at 86.34 %, with the lowest accuracy at 75.19 % *(Schizosaccharomyces pombe*).

**Table 3.** Summary of experimental evidences found in STRING [39]. <u>organism</u>: latin name for eight model organisms sorted alphabetically; <u>NpredPPIs</u>: number of PPIs of 1% ranked predictions; <u>NEvidence</u>: number of PPIs for which experimental evidences was found in at least on of the three databases used for training; <u>NcorrectEvidence</u>: number of PPIs with experimental evidence which were correctly classified by our approach; <u>Accuracy</u>: fraction of correct predictions within the predictions with experimental evidence.

| Organism | NpredPPIs | NEvidence | NcorrectEvidence | Accuracy |
| --- | --- | --- | --- | --- |
| *A. thaliana* | 250,000 | 1,138 | 1,138 | 100.00 % |
| *C. elegans* | 200,000 | 1,671 | 1,818 | 91.91 % |
| *D. melanogaster* | 100,000 | 1,763 | 2,049 | 86.04 % |
| *E. coli* | 40,000 | 1,807 | 2,196 | 82.29 % |
| *M. musculus* | 200,000 | 0 | 0 | - |
| *P. falciparum* | 50,000 | 0 | 0 | -% |
| *S. pombe* | 50,000 | 1,088 | 1,447 | 75.19 % |
| *R. norvegicus* | 200,000 | 0 | 0 | - |
| *Sum over all 8* | 1,090,000 | 7,467 | 8,648 | 86.34 % |

Fig. 3 illustrates the distribution of the experimental evidences found in STRING [39] plotted against their STRING scores. For *Arabidopsis thaliana* (Fig. 3, first row, first column), evidences in STRING were found for only positive predicted PPIs (1138 evidences). This results in about ≈ 70 % of the predictions having a STRING confidence score between 0.4 and 0.6, and the remaining ≈ 30 % having a high confidence score between 0.6 and 1.0. For *Caenorhabditis elegans* (Fig. 3, first row, second column) and *Escherichia coli* (Fig. 3, second row, first column), the accuracy of positive predicted PPIs found in STRING amounts to respectively 91.91 % and 82.29 %. Plotting the distribution of positive and negative predicted evidences found in STRING, both plots for *Caenorhabditis elegans* and *Escherichia coli* show similar distribution between positive and negative predicted PPI. In both cases, we found equal distribution of lower and higher STRING confidence score for both positive and negative predicted PPIs. In contrast, *Drosophila melanogaster* (Fig. 3, first row, third column) and *Schizosaccharomyces pombe* (Fig. 3, second row, second column) show a difference in distribution between positive and negative predicted PPIs. We observe a high percentage of STRING scores (below 0.5 for more than 80 % of the evidences) for negative PPIs, and a high percentage of high STRING scores (above 0.7 for 50 % of the evidences found). The negative predictions which were still found in STRING are likely to be false positive, as according to [39]: “A score of 0.5 would indicate that roughly every second interaction might be erroneous (i.e., a false positive).”

**Fig 3.**
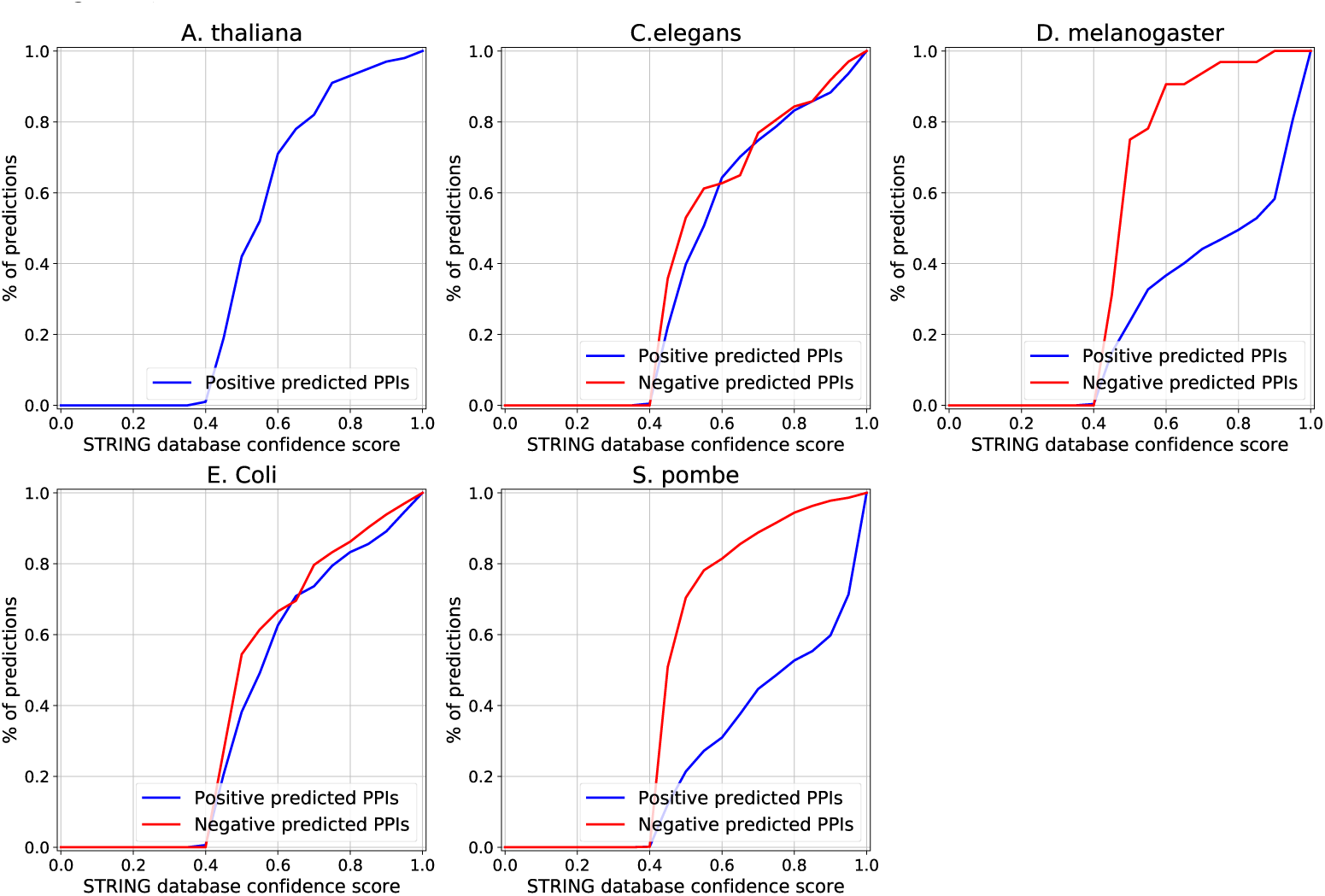
Percentages of predictions as a function of STRING [39] (confidence) score. The fractions of positive and negative predicted PPIs are each plotted against their STRING database confidence score. The plots show the plots for evidences for *Arabidopsis thaliana, Caenorhabditis elegans, Drosophila melanogaster, Escherichia coli* and *Schizosaccharomyces pombe*. For *Mus musculus* and *Rattus norvegicus*, no evidence was found.

### Insights from novel predictions

The majority of PPIs predicted by our models has not been reported in any of the three databases that we used at any level of reliablity (BioGRID, DIP, and IntAct). Column 4 of Table 2 (NpredPPI novel) summarizes the number of novel PPIs predicted for each organism; novel means that they differ from all experimentally known PPIs, including redundant and low-quality PPIs. Even if we assumed that only one in 20 of the positive predictions were right, these large numbers demonstrated that even for the best studied organisms, millions of PPIs without a close homolog from which interactions could be inferred remain unknown.

What can be stated about those newly predicted PPIs? While there is no answer for the millions, we investigated the most reliable 100 PPI predictions for *Escherichia coli* (note ‘only’ about 300k PPIs were predicted novel in *Escherichia coli*). 79 of these 100 PPIs were annotated to involve DNA-binding proteins. We are aware of very few DNA-binding proteins that do not bind to other proteins. Thus, the fact that DNA-binding proteins are involved in almost 80 % of all our top predictions of PPIs that were never seen before seemed at least encouraging. However, we did not find any clear evidence supporting any one of those 79 PPIs explicitly. 15 of the 100 top PPIs were annotated to involve repressing molecular binding. For example, *Escherichia coli* proteins P0ACP7 and P0ACQ0 were predicted with strong reliability (probability = 0.999977). Both proteins were classified as repressors by UniProt [42]. Transcriptional repression is an important aspect of gene regulation. As in most areas of molecular biology, studies of *Escherichia coli* have provided the model for subsequent investigations of transcription in different organisms, in particular in eukaryotic cells [43]. We were, therefore, surprised that some of our strongest predictions of PPIs never seen before involved *Escherichia coli* repressors. Again, we did not find any explicit experimental data to support or refute these 15 novel PPI predictions.

Further findings include Zinc finger (ZnF) domains, which are widely distributed in eukaryotic genomes. It has been estimated that around 1% of all genes encode proteins containing ZnFs and those proteins often contain multiple repeats of ZnFs [44]. Their functions are extraordinarily diverse and include DNA recognition, RNA packaging, transcriptional activation, regulation of apoptosis, protein folding and assembly, and lipid binding. Zinc finger structures are as diverse as their functions. In general, little is known about these protein–protein interactions [45]. We analysed the molecular function using Gene Ontology (GO, [46]). Interestingly, zinc ion binding is a molecular function which 81 of the top 5000 *Drosophila melanogaster* protein pairs of positive predicted PPIs have in common as well as 5 of the top 1000 *Caenorhabditis elegans* PPIs. However, protein pairs both being zinc ion binding in *Arabidopsis thaliana(* 181 of the top 5000) and in *Schizosaccharomyces pombe*(7 of the top 1000) are common functions of protein pairs highly unlikely to interact. Similar to our findings about *Escherichia coli*, we did not find any explicit experimental data to support or refute these interactions relating to zinc ion binding protein pairs.

### Limitation of performance evaluation

Several problems were in the way to providing a completely convincing comprehensive performance assessment. Specific to our problem were the rather small data sets of experimentally characterized PPIs: fewer than 6,000 non-redundant PPIs for all 8 organisms. In order to avoid severe problems from database bias, we had to focus on high-quality non-redundant PPIs [23]. As our profile-kernel based SVM requires at least 200 reliable PPIs, the number of acquired non-redundant PPIs reduced the set of organisms to only 8. The additional challenges were not specific to our work: it remains uncertain by more than an order of magnitude how many interactions are to be expected in an organism. Related to this: what is the fraction of positives (PPIs) to negatives (proteins that do not interact) is in a living cell? Yet another crucial problem is that positives are much more reliable than negatives. For molecular biology in general it is much more accurate to state that an event happens than to rule out that it does not. All these issues magnify each other to render even the most careful performance estimates to become speculative approximations at best. Many authors use ROC-curves that relate the number of true positives (correctly predicted PPIs) to that of false positives (PPIs predicted but not observed). These plots depend heavily on the negatives in particular on the ratio of positives-to-negatives. Given that the truth for this number remains uncertain even within an order of magnitude, we decided to focus on curves that show precision-vs-recall, i.e. only values directly related to the observed PPIs. Although one of the axes still is strongly influenced by the assumption that ‘not observed’ means ‘not interacting’. AUC, the area under the ROC-curve, is another simple and popular score for performance evaluation. Given the argument against ROC-curve, we might still vary this and compile an analogous area under the precision-recall curve. However, such a value would constitute another major problem: arguably, most users of prediction methods are most interested in the most reliable predictions. In other words, when predicting whether protein X interacts with any other human protein, the N-strongest predictions (for some N might be 1 for others 1000) matter more than all 20k scores against all 20k human proteins. But those 20k-N would exactly dominate the AUC-type performance measures.

### Database of predictions

Table 2 summarizes the results of the full interactome predictions. We only predicted PPIs which are dissimilar to proteins in our positive training set (Table 2, column NprotPred). Most proteins of the reference proteomes were dissimilar (Table 2: difference between columns Nprot, number of proteins, and NprotPred, number of predicted proteins). Overall, the eight new methods predicted PPIs for most of all possible pairs of proteins in an organism, i.e. at least 73 % of all possible pairs (only exception: *Escherichia coli* and *Drosophila melanogaster*). Even after excluding all proteins previously reported in low-quality or redundant PPIs from the set of predicted PPIs, millions of predicted PPIs remained (Table 2, column NpredPPI novel). Due to our large mistake in the prediction of all PPIs proposed by the model at the default threshold, the ProfPPIdb resource only reported the most reliably predicted, non-redundant predictions (top ~ 10 % of all predicted PPIs) as novel PPIs (Table 2, column NpredPPI novel). For most of the 8 model organisms, this subset excludes most proteins in the organism (Table 2, numbers in column NprotPred more than twice those in column Nprot ProfPPIdb). The exceptions were Plasmodium falciparum, Schizosaccharomyces pombe and Drosophila melanogaster for which PPI predictions remained for almost all proteins with predictions (Table 2, column NprotPred) after the application of these filters (Table 2, column Nprot ProfPPIdb). Hence, although our resource adds over one million newly predicted PPIs (sum over 8 rows of column NpredPPI ProfPPIdb in Table 2: 1,090,000 PPIs), many proteins in those organisms remain without annotation and without predictions.

## Conclusions

We applied the concept of profile-kernel SVMs for the prediction of physical protein-protein interactions (PPIs), i.e. we leverage information available for all proteins for which the sequence is known. The profile-kernel SVM-based methods appeared to achieve state-of-the-art performance for sequence-based PPI predictions. In fact, for most model organisms, the predictions were not inferior to those for human for which we had most experimental data and developed our initial approach. We put the most reliable predictions into a freely available database where users can access predictions for all proteins in the entire proteomes of eight different organisms (eukaryotes and prokaryotes, multi-cellular and single cellular, animals and plants, mammals, fly and worm).

## Acknowledgments

Linh Tran is now at Imperial College London, but did this work earlier as a Technical University of Munich student. Thanks primarily to Tim Karl, but also to Guy Yachdav (all TUM) for invaluable help with hardware and software; to Inga Weise (TUM) for support with many other aspects of this work; to Jonas Reeb and Lothar Richter (both TUM) for help in supervising some of the work. This work was supported by a grant from the Alexander von Humboldt foundation through the German Ministry for Research and Education (BMBF: Bundesministerium fuer Bildung und Forschung). Last, not least, thanks to all those who deposit their experimental data in public databases, and to those who maintain these databases. In particular thanks to the BIND team headed by Gary Bader and Christopher Hogue, the DIP team around David Eisenberg, Ioannis Xenarios, and to the IntAct team co-ordinated by Henning Hermjakob.

## A Supporting Information

### A.1 Cross validation results

#### Similar levels of training and holdout performances

Machine learning applications often reach very different levels of performance for the training and the testing set. We did not observe this for the organisms for which we could compile comprehensive cross-validation results (Fig. 4: difference between black line and colored points). Most similar were the results for mouse *(Mus musculus:* Fig. 4 E). For *Escherichia coli* (Fig. 4 A), *Caenorhabditis elegans* (worm, Fig. 4 B), and *Drosophila melanogaster* (fruit fly, Fig. 4 C), training and testing were less similar for high recall, i. e. for the most reliable predictions. Most unusual were the results for *Drosophila melanogaster* (Fig. 4 C) and *Escherichia coli* (Fig. 4 A), for which test performance was even higher than training performance for a substantial fraction of highly reliable predictions (toward left, i.e. low recall in Fig. 4 A, and Fig. 4 C the black curves are above the dots). For *Arabidopsis thaliana* (water-cress, Fig. 4 D) testing performance was better than training throughout the entire ROC-like curve. Typically, there is only one explanation for such unexpected findings: points for which testing is better than training provide estimates for the resolution of our performance estimates. This reality was captured well by the estimates for standard errors: within one standard error, training and testing were identical for all organisms.

**Fig 4.**
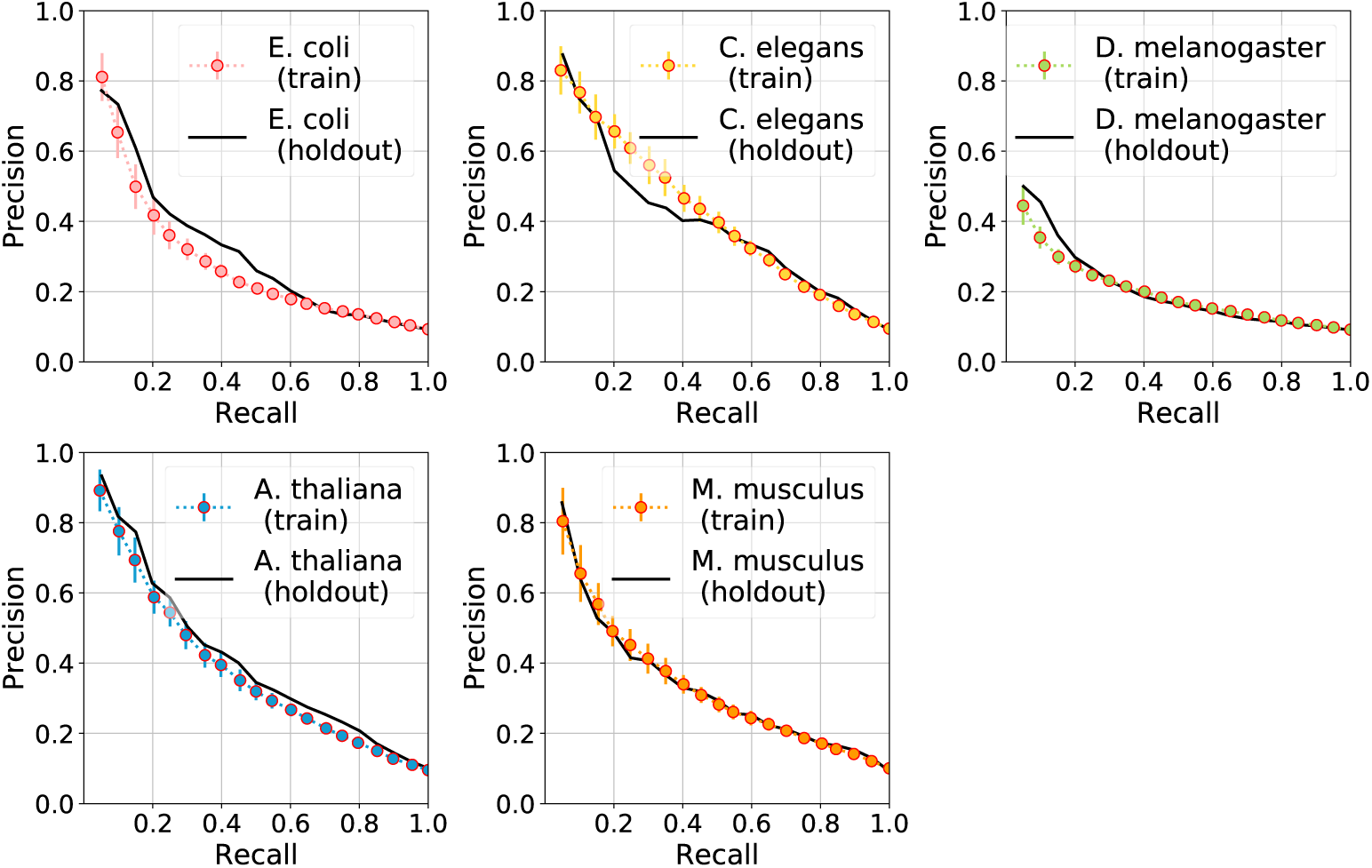
Cross-validation and holdout performance results for organisms with train data size > 500 PPIs. The y-axes give precision (number of PPIs correctly predicted at threshold), the x-axes the recall (number of experimental interactions predicted at that threshold). Bars give the standard deviation; negatives were sampled at a rate of 10:1 (ten negatives for one positive). Each subplot is referred as follows: A (*Escherichia coli)*, B *(Caenorhabditis elegans)*, C *(Drosophila melanogaster)*, D *(Arabidopsis thaliana)*, E *(Mus musculus)*.

#### Hyperparameter optimization for *Escherichia coli*

Our most important objective when applying machine learning typically is to reduce the risk of over-optimization, i.e. to optimize generalization instead of apparent performance as usually over-estimated by standard cross-validation. Therefore, we trained each organism model with the same set of hyperparameters (k-mer = 5 and *σ* = 11). This standard choice yielded the best performance for almost all organisms. One exception was Escherichia coli. For the choice k-mer = 3 and *σ* = 4, the cross-validation precision-recall values exceeded those for all other hyperparameter combinations (Fig. 5 A). This top choice for *Escherichia coli* reached higher performance than the human-specific model in the realm of low recall (Fig. 5 B). This choice for *Escherichia coli* also results in high performance for the holdout set of E.coli which exceeds the test performance of *Homo sapiens* from [23] especially in the realm of low recall (Fig. 5C).

**Fig 5.**
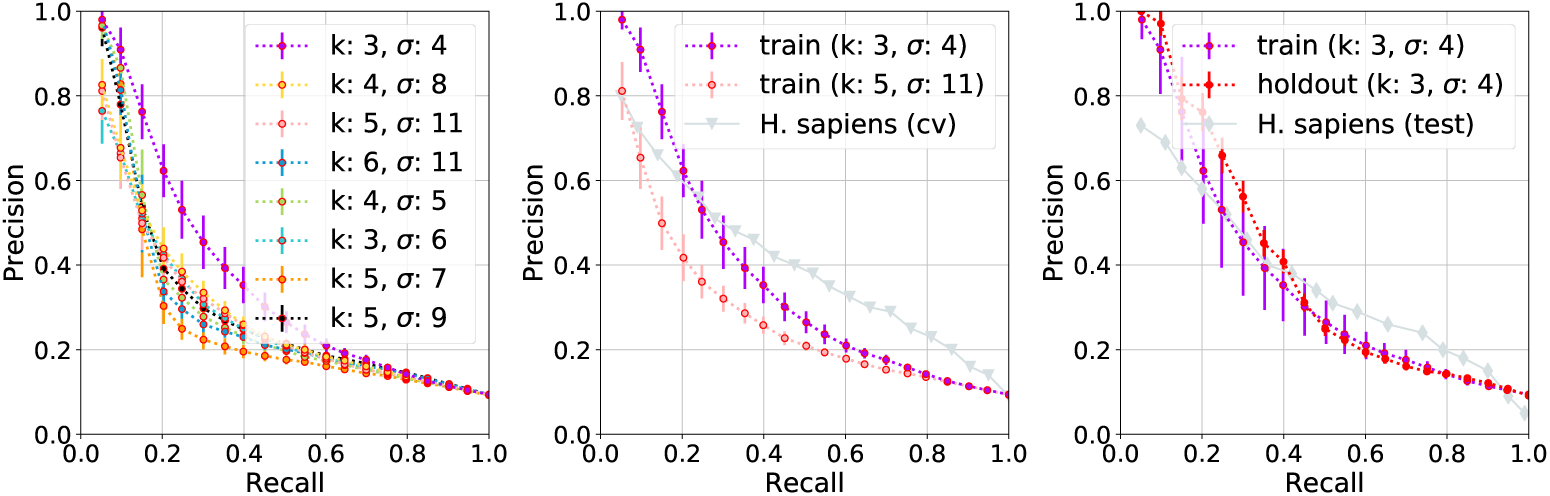
Cross-validation and hold-out performance results of *Escherichia coli*. **<u>Panel (A)</u>**: Precision-recall curve for cross-validation in *Escherichia coliw*ith different optimization hyperparameters. All results in the paper were reported for the version *k = 5/σ* = 11 which clearly was not best for *Escherichia coli*, instead the combination *k* = 3/σ = 4 yielded the best performance (purple**). Panel (B)**: Comparison of cross-validation hyperparameter combinations *k* = 3/*σ* = 4 (best) with *k* = 5/*σ* = 11 (default) and cross validation of human from earlier publication [23]. **Panel (C):** Cross-validation and hold-out results of hyperparameter combination *k* = 3/*σ* = 4 (best) compared with test results for human [23].

### A.2 Evaluation of novel predictions

We used **BioGRID** [25], **DIP** [26], and **IntAct** [27] (Uniprot uses quality-filtered subset of binary interactions automatically derived from the IntAct database) for large-scale evaluation of our novel predictions. Although we used BioGRID [25], DIP [26], and IntAct [27] as the base for our organism-specific models, it was only a small subset of the databases’ PPIs actually used for training our models.

The PPIs published on our online service only include PPIs which have not any experimental evidence from any of these three databases. In order to perform an evaluation of the quality of the predictions, we used the top 1 % of all predictions (ranked according to our confidence measure) which were not included in the training set. We compared these top predictions against all experimental from BioGRID [25], DIP [26], and IntAct [27]. The findings of experimental evidences is listed in Table 4. As Table 4 shows, except for Mus musculus and Rattus norvegicus for which none or only falsely predicted PPIs was found, we found between 60 and 170 PPIs with experimental evidence for each organism. The accuracy of the evidences correctly predicted is at least over 75 %, with half of all investigated organisms having accuracies of over 90 %.

**Table 4.** Summary of experimental evidences found in BioGRID [25], DIP [26], and IntAct [27]. <u>Organism</u>: latin name for eight model organisms sorted alphabetically; <u>NpredPPIs</u>:Number of PPIs of 1% ranked predictions; <u>NEvidence</u>: number of PPIs for which experimental evidences was found in at least on of the three databases used for training; <u>NcorrectEvidence</u>: number of PPIs with experimental evidence which were correctly classified by our approach; <u>Accuracy</u>: The fraction of correct predictions within the predictions with experimental evidence.

| Organism | NpredPPIs | NEvidence | NcorrectEvidence | Accuracy |
| --- | --- | --- | --- | --- |
| *A. thaliana* | 2,064,410 | 62 | 60 | 96.77 % |
| *C. elegans* | 1,421,719 | 67 | 69 | 97.1 % |
| *D. melanogaster* | 319,160 | 152 | 197 | 77.16 % |
| *E. coli* | 27,296 | 82 | 90 | 91.11% |
| *M. musculus* | 1,313,253 | 0 | 0 | - |
| *P. falciparum* | 90,418 | 143 | 174 | 82.18% |
| *S. pombe* | 83,497 | 166 | 177 | 93.79% |
| *R. norvegicus* | 2,119,225 | 3 | 0 | 0.00 |
| Sum over all 8 | 6,125,724 | 772 | 670 | 86.79 % |

Looking closer at the distribution of the evidences in terms of average, we found three cases which we show in Figure 6. With E. coli (Figure 6 a), we observe a high percentage of lower average expert knowledge scores (below 4 for almost 80 % of the evidences) for negative PPIs, and a high percentage of high average expert knowlege scores (greater or equal 6 for 60 % of the evidences found). This shows that for E. coli our model succeeds in predicting PPIs correctly which also has experimental evidences with high average expert knowledge scores. However, for organisms A. thaliana (Figure 6 b), C. elegans, D.melanogaster, P. falciparum and S. Pombe we witness PPI curves which almost overlaps. This indicate a similar distribution of knowledge expert scores for both positive and negative PPIs. This also is a consequence of the lack of high quality annotations present in the databases. The third case of distribution that we observed is with M. musculus, for which we only found three experimental evidences and none of them are correctly predicted by our approach. Nevertheless, the experimental evidences are highly doubtful as their methods each score only 1 in the expert knowledge scoring system.

**Fig 6.**
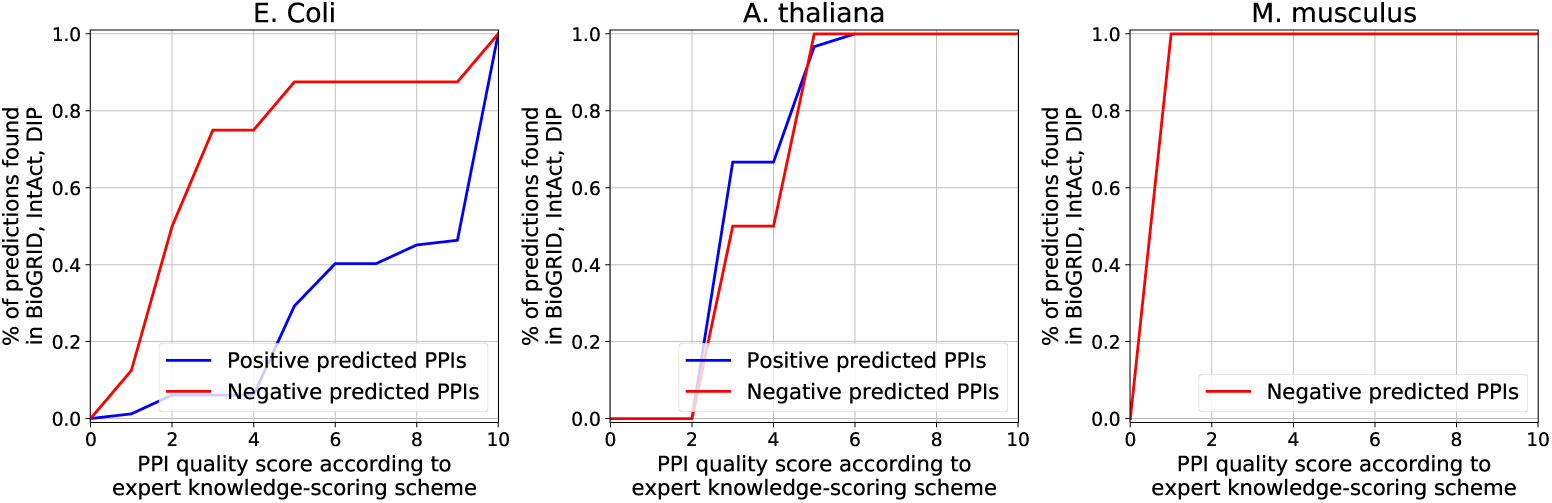
Percentages of predictions as a function of PPI quality score according to expert knowledge scoring scheme [29]. This scoring scheme was also used in the manuscript to obtain high-quality PPIs for training. The positive and negative PPIs presented in these plots are findings of experimental evidences found in BioGRID [25], DIP [26], and IntAct [27].

